# Apolipoprotein A-I Mimetic 4F Peptide Generates Amyloid Cytotoxins by Forming Hetero-oligomers with β-amyloid

**DOI:** 10.1101/722983

**Authors:** Bikash Ranjan Sahoo, Michael E. Bekier, Zichen Liu, Vojc Kocman, Andrea K. Stoddard, G. M. Anantharamaiah, James Nowick, Carol A. Fierke, Yanzhuang Wang, Ayyalusamy Ramamoorthy

## Abstract

Apolipoproteins are involved in pathological conditions of Alzheimer’s disease (AD), truncated apolipoprotein fragments and β-amyloid (Aβ) peptides coexist as neurotoxic heteromers within the plaques. Therefore, it is important to investigate these complexes at the molecular level to better understand their properties and roles in the pathology of AD. Here, we present a mechanistic insight into such heteromerization using a structurally homologue apolipoprotein fragment of apoA-I (4F) complexed with Aβ(M1-42) and characterize their toxicity. The 4F peptide slows down the aggregation kinetics of Aβ(M1-42) by constraining its structural plasticity. NMR and CD experiments identified 4F-Aβ(M1-42) heteromers as being comprised of unstructured Aβ(M1-42) and helical 4F. A uniform ≈2-fold reduction in Aβ42 ^15^N/^1^H NMR signal intensities with no observable chemical shift perturbation indicated the formation of a large complex, which was further confirmed by diffusion NMR experiments. Microsecond scale atomistic molecular dynamics simulations showed that 4F interaction with Aβ(M1-42) is electrostatically driven and induces unfolding of Aβ(M1-42). Neurotoxicity profiling of Aβ(M1-42) complexed with 4F confirms a significant reduction in cell-viability and neurite growth. The molecular architecture of heteromerization between 4F and Aβ(M1-42) discovered in this study provides evidence towards our understanding of the role of apolipoproteins or their truncated fragments in exacerbating AD pathology.

## Introduction

Aberrant protein folding and aggregation are associated with a variety of neurodegenerative disorders including Alzheimer’s disease (AD), Parkinson’s disease and prion diseases^1–3^ and non-neuropathic disease like type-2 diabetes (T2D). The progression of these diseases is linked with the accumulation of insoluble fibrillary aggregates (also called amyloid-plaques) from water-soluble protein monomers such as β-amyloid (Aβ) cleaved from the amyloid-precursor protein (APP), which has been intensely investigated to support the amyloid hypothesis cascade. Although the molecular basis of amyloid-deposition is not fully understood, several studies have identified the formation of water-soluble Aβ peptide oligomers from monomers prior to the formation of metastable amyloid fibers. Interestingly, an increasing amount of evidence has recently shown the association of these cytotoxic oligomers (also called cytotoxins) to human neurons or pancreatic β-cells.^4^ The presence of oligomers surrounding amyloid plaques has also been identified as a distinguishable morphological hallmark in patients with dementia as compared to healthy people with a similar plaque morphology but not dementia.^5^ Although the characterization of such oligomers (namely amylospheroids, protofibrils, paranuclei, globulomers etc.) remains challenging due to their morphological heterogeneity, their relatively high neurotoxicity as compared to monomers or amyloid fibers directs researchers to target these species for therapeutic developments.

While there is significant interest in fully characterizing Aβ oligomers due to their roles in the pathogenesis of AD,^6^ several studies have shown the co-occurrence of other cellular cofactors including proteins^7^ such as apolipoprotein fragments in the formation of hetero-oligomers.^8^ The 35-kDa apolipoprotein-E (apoE) and proteolytically cleaved apoE fragments are associated with Aβ and have recently been identified from the brain of an AD patient.^8–10^ Accumulation of apoE-Aβ complex has been identified in AD cortical synapses from human brain specimens.^11^ apoE is involved in the modulation of Aβ metabolism, aggregation and clearance,^12,13^ a recent in vitro study showed that apoE3, apoE4 and different fragments of apoE4 derived from N- or C-terminus substantially delay Aβ(1-42) aggregation kinetics.^14^ Other neuroprotective apolipoproteins such as apoA-I and apo-J have also been observed in AD brains and heteromerization of apoA-I/Aβ has been suggested to play a neuroprotective role.^15–19^ provide strong evidence that truncated apolipoprotein fragments and their interactions with Aβ play critical roles in AD progression.

Several approaches have been used to directly modulate amyloid aggregation and oligomerization pathways as possible AD treatments including binding of small-molecule inhibitors, peptides, polymers, cellular co-factors like ions, cell-membrane, enzymes, and chaperones.^20–24^ Other indirect approaches such as enhancing Aβ clearance pathways to control AD progression have also been reported.^25^ One such approach is the enhancement of apoE secretion that promotes Aβ cellular trafficking and degradation by using apolipoprotein mimetic peptides.^26^ Nevertheless, apoE antagonist has been designed to block the interaction between APP and apoE that enhances Aβ production.^27^ Recently, an apoA-I mimetic peptide (4F) has been shown to directly inhibit Aβ(1-40) aggregation ^28^ and was also identified to upregulate apoE secretion to promote Aβ’s clearance.^26^ Moreover, the 4F peptide has also been shown to have an excellent medical application for several other human diseases.^29–32^ These findings urge a need for deeper understanding of their biomedical application in amyloidosis especially in AD progression.

Several studies have used short peptide inhibitors/modulators for AD treatment; however, none of them have been successful in clinical trials indicating our poor understanding of the molecular basis of the disease progression. Furthermore, blocking Aβ aggregation may even increase the risk of generating cytotoxic Aβ oligomers or other intermediates.^33–35^ Alternative approaches such as promotion of quick fibrillation of Aβ by chemical modulators to suppress the generation of cytotoxic intermediates have recently been reported ^36–38^. Together with the recent discovery of apoE-fragment/Aβ heteromer in AD brains,^8^ this raised a concern that the 4F inhibition of Aβ may possibly generate cytotoxic Aβ species.^28^ Similarly, a previous in vivo study showed the apoE-Aβ complex reduces the level of soluble species and increases the Aβ oligomer levels that are known to be neurotoxic.^39^ The binding of apoE to Aβ oligomers at substoichiometric and to fibers at high concentration has also been demonstrated in vitro indicating their crucial involvement during amyloid fibrillation.^40^ Therefore, there is an urgent need to investigate the mechanism of heteromerization between apolipoprotein fragments and Aβ in correlation with their toxicity and disease progression. The insights gained from such mechanisms could enable the design of better therapeutic strategies to detoxify these cytotoxins.

Herein, we investigated the interaction between the apolipoprotein peptide fragment 4F and Aβ(M1-42). Our results reveal a substantial binding between 4F and Aβ(M1-42) as recently observed for Aβ_1-40_. A structural model showing the formation of hetero-oligomers is developed using a combination of biophysical methods including NMR and multi-microseconds all-atom and coarse-grained molecular dynamics (MD) simulations. Pathological phenotype characterization of the isolated 4F-Aβ(M1-42) hetero-oligomer shows an elevated cellular toxicity as compared to Aβ(M1-42) aggregates.

## Results and Discussion

### 4F peptide retards Aβ(M1-42) fibrillation

To investigate the involvement of apolipoproteins and their fragments in the pathology of AD and T2D, we selected apoA-I mimetic 4F peptide that share structural homologs with C-terminal apoE fragments and apoA-I.^8,41^ The effect of 4F peptide on the aggregation kinetics of Aβ(M1-42) was first monitored using thioflavin-T (ThT) based fluorescence assay. In the absence of 4F peptide, Aβ(M1-42) (5 µM) showed a very short lag-time (< 2 hour) of aggregation (Fig. 1a). On the other hand, a significant difference in Aβ(M1-42) aggregation kinetics was observed in the presence of equimolar 4F. The lag-time of Aβ(M1-42) fibrillation was increased by a factor of ≈24 to keep the amount of fibril nearly the same as observed in the absence of 4F (Fig. 1a). With further increase in 4F concentration (i.e. at 25 or 50 µM), a substantial fluorescence quenching and retardation in aggregation kinetics (up to 48 hours) were observed for Aβ(M1-42) which correlate to our previous observations for Aβ (1-40).^28^ Morphological analysis of the reaction end product by transmission electron microscopy (TEM) displayed a mixture of Aβ(M1-42) fibers and spherical oligomers in samples without 4F (Fig. 1b). In presence of equimolar 4F, short and comparatively thick fibers of Aβ(M1-42) were identified that are morphologically different from 4F or Aβ(M1-42) alone in solution (Fig. 1b-d). At 50 µM 4F, Aβ(M1-42) showed comparatively very small amount of fiber (straight fiber morphology) with few globular oligomers that correlate the ThT observation (Fig. S1). Taken together, the fluorescence and TEM results indicate that the 4F peptide has a more pronounced effect on slowing down Aβ(M1-42)’s aggregation by generating morphologically distinct species.

**Fig. 1.**
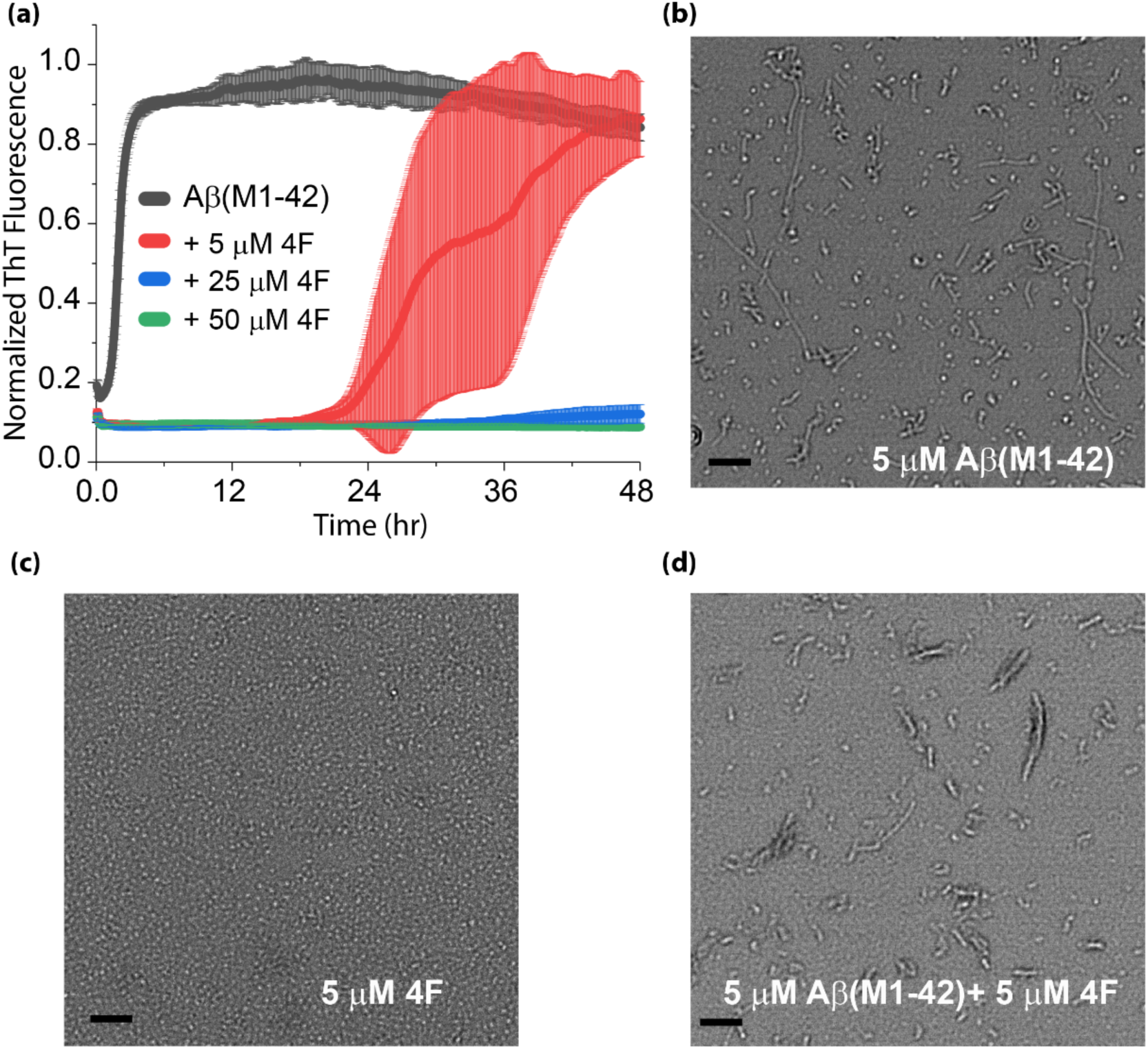
(a) Time course aggregation kinetics of 5 µM Aβ(M1-42) monitored using ThT fluorescence in presence and absence of 4F peptide at variable concentration as indicated in different colors. The average ThT curve of the samples in triplicate are shown in a solid line and standard errors are highlighted in color. (b-d) TEM images of 5 µM Aβ(M1-42) with/without equimolar 4F peptide. Samples for TEM experiments were taken from the 96-wells plate used for ThT fluorescence experiments but had no ThT dye. The scale bar is 200 nm.

### 4F peptide stabilizes the aqueous conformational state of Aβ(M1-42)

Next, we investigated the inhibitory mechanism of the 4F peptide by monitoring the conformational change in Aβ(M1-42). Freshly prepared Aβ(M1-42) monomers incubated with or without 4F peptides were analyzed for several days by circular dichroism (CD) and ^1^H NMR experiments. CD results showed an initial random-coil rich and partially folded structure for 20 µM of Aβ(M1-42) and 4F, respectively. Notably, when both peptides were mixed at equimolar ratio (20 µM), a prominent α-helical conformation characterized by negative peaks located between ≈208 and ≈222 nm was identified (Fig. 2a) with a substantial increase (≈ 3-fold as compared to 4F alone) in the molar ellipticity (Θ). Time-lapse CD measurements further identified the stability of the α-helical structure in 4F-Aβ(M1-42) mixed solution with no change in ‘Θ’ and CD minima for a duration of 2 days (Fig. 2a). Thereafter, a decrease in ‘Θ’ was observed with a negligible change in CD minima possibly due to amyloid fibrillation. Although CD showed a distinct conformational transition in the 4F-Aβ(M1-42) complex, it was not possible to ascertain which of the two different peptides has contributed to the induction of the experimentally observed α-helical secondary structure.

**Fig. 2.**
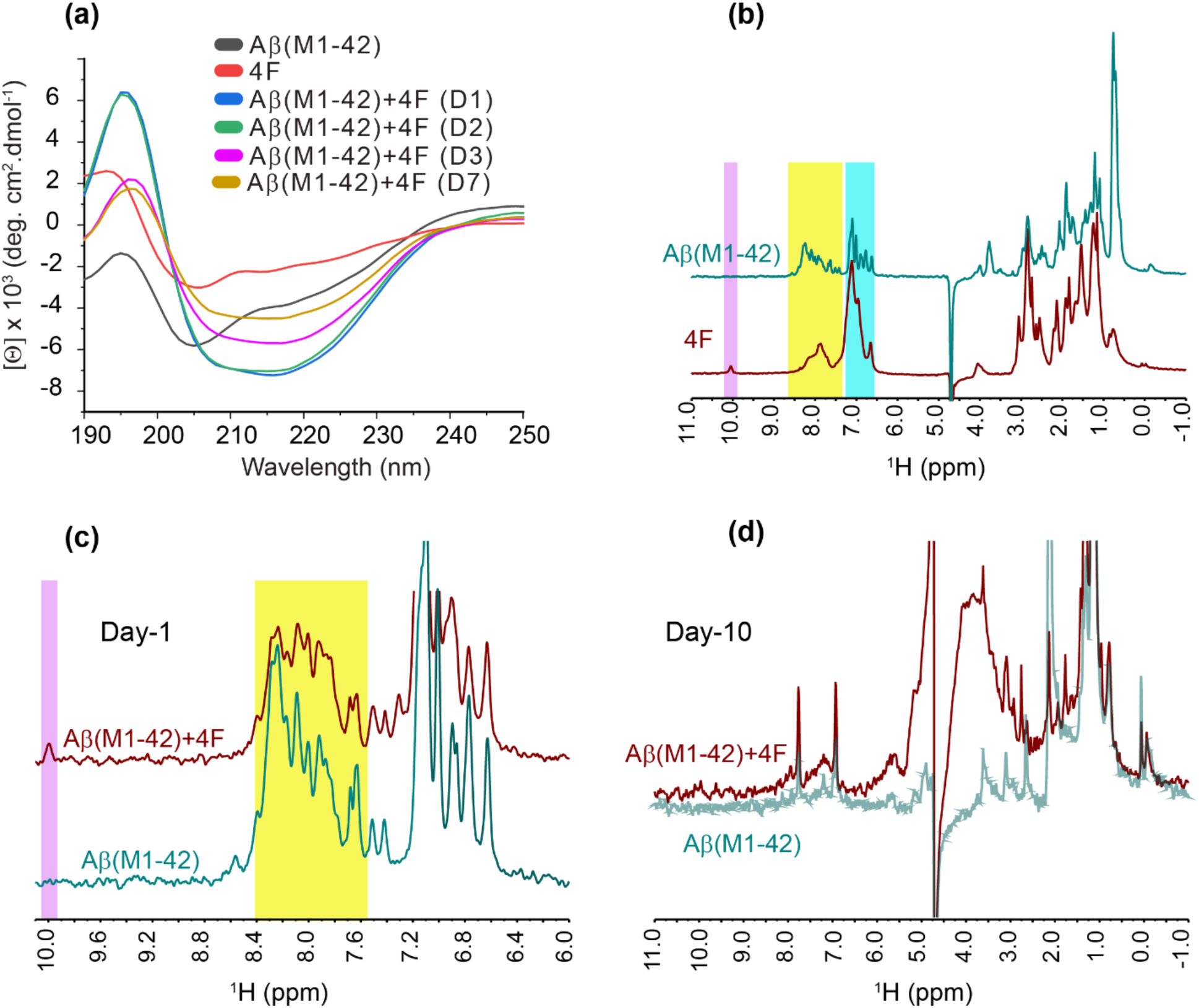
(a) Time-lapse measurement of conformational change in 20 µM Aβ (M1-42) mixed with equimolar 4F peptide using CD spectroscopy. The samples were incubated at room temperature and CD measurements were carried out until day-7 (denoted as D7). (b) ^1^H NMR spectra of 20 µM Aβ (M1-42) and 4F peptides. The distinguished amide-proton (N-H) and 4F peptide’s Trp indole NH proton peaks are highlighted. (c) ^1^H NMR spectra illustrate the changes in amide proton resonances in 20 µM Aβ (M1-42) mixed with an equimolar 4F peptide. The yellow region highlighted a small change in the Aβ (M1-42) spectrum pattern mixed with 4F. (d) ^1^H spectra of NMR samples used for (c) after 10 days incubation under gentle shaking at 37 °C. All NMR spectra were recorded on a 500 MHz NMR spectrometer at 25 °C.

To explore the conformational states exhibited by the two different peptides in the 4F-Aβ(M1-42) mixed solution, we next carried out NMR experiments. The fingerprint amide N-H region in proton NMR spectra was used to monitor the conformational state of individual peptides.^42^ As illustrated in Fig. 2b, the amide protons showed a distinct chemical shift dispersion for Aβ(M1-42) and 4F. The Aβ(M1-42) peptide exhibited sharp amide proton peaks in the ≈7.5 to 8.5 ppm region; whereas, the 4F peptide showed broad peaks in that region and a distinct isolated peak for the indole ring N-H proton at ≈10 ppm. The ≈7.5 to 8.5 ppm region of the NMR spectrum of Aβ(M1-42) mixed with equimolar 4F showed an NMR peak pattern similar to that of Aβ(M1-42) alone in solution (Fig. 2c). Since most of the fingerprint region (7.5 to 8.5 ppm) for Aβ(M1-42) remains unchanged after the addition of 4F, we hypothesize that the random-coil rich Aβ(M1-42) conformation remained unaffected when bound to 4F.^42^ Aβ(M1-42) peptide alone (with no additives) was incubated for several days to allow for peptide aggregation and the progress was monitored by proton NMR (Fig. 2c and d). Proton NMR spectra recorded on day-10 presented substantial broadening and intensity decrease for most of the peaks (Fig.2d) as compared to day-1 (Fig. 2c), indicating Aβ(M1-42)’s fibrillation both in absence and presence of 4F. Interestingly, proton NMR peaks from 4F including the tryptophan indole ring N-H proton (at ∼10 ppm as seen in Fig.2b) were absent, and a similar amide proton NMR peak pattern was observed for both samples (Fig. 2d). This indicated a likely complexation of 4F peptide molecules with Aβ(M1-42) fibers. Such aggregates in general exhibit line broadening and are beyond detection by solution NMR. The stability of the complex was further investigated by washing these large aggregates in 2M NaCl (see Methods). Tryptophan fluorescence of the filtrate sample, with fluorescence emission at ≈458 nm when excited at 295 nm, confirmed the presence of the 4F peptide indicating the disintegration of 4F from Aβ(M1-42) fibers (Fig. S2a). However, it did not rule out the possible presence of any 4F peptide in Aβ(M1-42) fiber after repeated washing using 2M NaCl. To explore this, the filtered Aβ(M1-42) fibers were incubated overnight with 2% deuterated SDS followed by 30 minutes heating at 80 °C (see Methods) and used to acquire proton NMR spectra which depicted no characteristic peaks corresponding to 4F; specifically, the SDS treated sample did not show the fingerprint tryptophan indole N-H proton (at ∼10 ppm) indicating the absence of 4F peptide (Fig. S2b) in filtered Aβ(M1-42) fibers.

### Secondary structure characterization of Aβ(M1-42) bound to 4F using NMR

We further investigated the secondary structure of Aβ(M1-42) bound to 4F in the heteromer state using 2D ^15^N/^1^H SOFAST-HMQC experiments. For this, we employed an approach to first monitor Aβ(M1-42)’s the overall secondary structure in two different solvent environments that give rise to a random-coil rich and a partially folded α-helix structure. 2D HMQC pattern and CD spectrum observed for Aβ(M1-42) in 10 mM sodium phosphate buffer resembled that of a random-coil like conformation reported previously (Fig. 3a).^43,44^ In the presence of 10% deuterated 2,2,2-Trifluoroethanol (TFE), a significant change in Aβ(M1-42) HMQC pattern and chemical shift was observed (Fig. 3b). Remarkably, a majority of Aβ(M1-42) ^15^N/^1^H peaks were found to depict upfield chemical shifts in ^1^H and ^15^N dimensions likely indicating Aβ(M1-42)’s conformational alteration (Fig. 3b). The upfield H-N chemical shifts in HMQC spectrum in presence of 10% TFE also indicated the presence of an α-helical Aβ(M1-42) as compared to its aqueous structure (Fig. 3a), and such chemical shift changes have been observed previously for several other proteins during folding.^45^ Further CD experiments on Aβ(M1-42) confirmed induction of an α-helix in Aβ (M1-42) in presence of 10% TFE (Fig. S3), The random-coil rich Aβ(M1-42) characterized by a negative peak at ≈200 nm showed an increase in ‘Θ’ and shift in CD minima that resemble an α-helical conformation (Fig. 3b). Induction of an α-helical conformation by TFE has also been observed for several short peptides including Aβ and thus correlate with the NMR observations mentioned above. Notably, our comparative study of Aβ(M1-42) conformation in presence of 10% TFE and 4F showed a very distinct HMQC pattern (Fig. 3a and b). Although a similar CD spectrum was obtained for Aβ(M1-42) in presence of 4F or 10% TFE (Figs. 2a and S3), a negligible ^15^N/^1^H chemical shift change for Aβ(M1-42) in presence of 4F and a substantial change in the presence of 10% TFE clearly indicated the existence of a random-coil rich Aβ(M1-42) bound to 4F (Fig. 3c). A comparison of the average chemical shift perturbation (Δδ_HN_) showed a majority of the Aβ(M1-42) residues have a chemical shift perturbation above the cut-off value in presence of 10% TFE; on the other hand, in presence of equimolar 4F the Δδ_HN_ value is negligible (Fig. 3c). Taken together, the NMR and CD results confirmed the strongly binding of 4F to Aβ(M1-42) to restrains Aβ(M1-42)’s conformational alteration, thereby indicating the mechanism of retardation of Aβ(M1-42)’s amyloid aggregation.

**Fig. 3.**
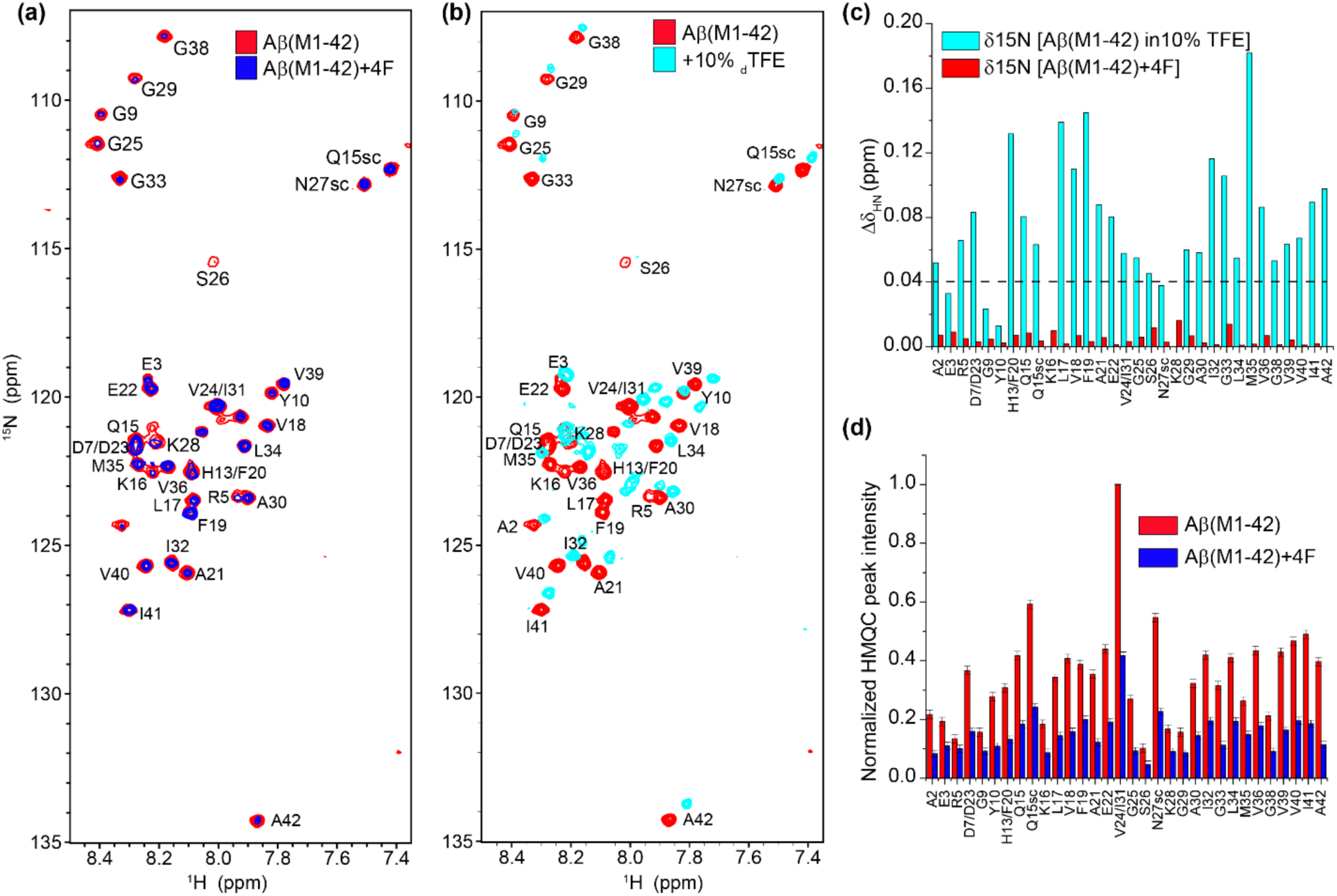
(a) Superimposed ^15^N/^1^H SOFAST-HMQC spectra of 20 µM uniformly-labelled (^13^C/^15^N) Aβ (M1-42) in 10 mM sodium phosphate, pH 7.4 mixed with (blue) or without (red) equimolar unlabeled 4F peptide. (b) ^15^N/^1^H SOFAST-HMQC spectra illustrating a conformation change in 20 µM Aβ (M1-42) in aqueous buffer (10 mM sodium phosphate, pH 7.4; in red) and buffer containing 10% deuterated TFE (in cyan). (c) The average chemical shift perturbation (Δδ_HN_) calculated from ^15^N/^1^H SOFAST-HMQC spectra in presence of equimolar 4F (Fig. 3A; in red) and 10% deuterated-TFE (Fig. 3b; in cyan) are plotted as a function of residue number. The dashed line in (c) indicates the average chemical shift perturbation observed for Aβ (M1-42) residues in 10% deuterated-TFE. (d) Normalized SOFAST-HMQC NMR peak intensities derived from Fig. 3a. Error bars represent standard error. NMR spectra were obtained NMR spectrometer at 25 °C.

### Atomistic insights into 4F and Aβ(M1-42) complex formation

The binding interactions and complex formation between 4F and Aβ(M1-42) at the atomic level was next investigated using 2D ^15^N/^1^H SOFAST-HMQC NMR experiments. 20 µM uniformly ^13^C/^15^N labeled Aβ(M1-42) in presence of equimolar 4F peptide exhibited a substantial reduction in the overall intensity for most of the residues in the 2D ^15^N/^1^H SOFAST-HMQC spectrum (Fig. 3). Aβ(M1-42)Aβ(M1-42)Notably, no significant difference in the chemical shifts of Aβ(M1-42) residues was observed indicating a negligible or no change in the Aβ(M1-42) conformation when bound to 4F (Fig. 3c). This observation is in good agreement with the above described CD and ^1^H NMR results Aβ(M1-42) shown in Fig. 2. On the other hand, the ^13^C/^1^H correlation spectrum of Aβ(M1-42) revealed the disappearance of resonance from the aromatic side chains of Phe and Tyr when mixed with equimolar 4F (Fig. S4). This observation indicates hydrophobic π–π packing could be the driving force for the complex formation between Aβ(M1-42) and 4F peptides. The uniform reduction in the NMR signal intensities in the 2D ^15^N/^1^H SOFAST-HMQC most likely due to a reduction in Aβ(M1-42)the mobility of Aβ(M1-42) residues when complexed with 4F which enhances the spin-spin relaxation and therefore broadens the observed NMR resonances. To further confirm this observation, we determined the diffusion rates of 4F mixed with and without equimolar Aβ(M1-42) using diffusion ordered NMR spectroscopy (DOSY). 20 µM 4F peptide in 100% D_2_O showed a diffusion constant of ≈ 9.65 m^2^s^-1^, whereas Aβ(M1-42) diffused faster with a diffusion rate constant of ≈8.93 m^2^s^-1^ (Fig. 4). Unlike the 4F peptide, the diffusion pattern observed for Aβ(M1-42) was found to be inhomogeneous due to the inherent nature of its aggregation during the NMR data acquisition (≈10 hours) (Figs. 4 and S5). Particularly, the 4F-Aβ(M1-42) mixed sample showed a diffusion pattern like that of the 4F indicating the presence of small-sized water soluble hetero-oligomers.

**Fig. 4.**
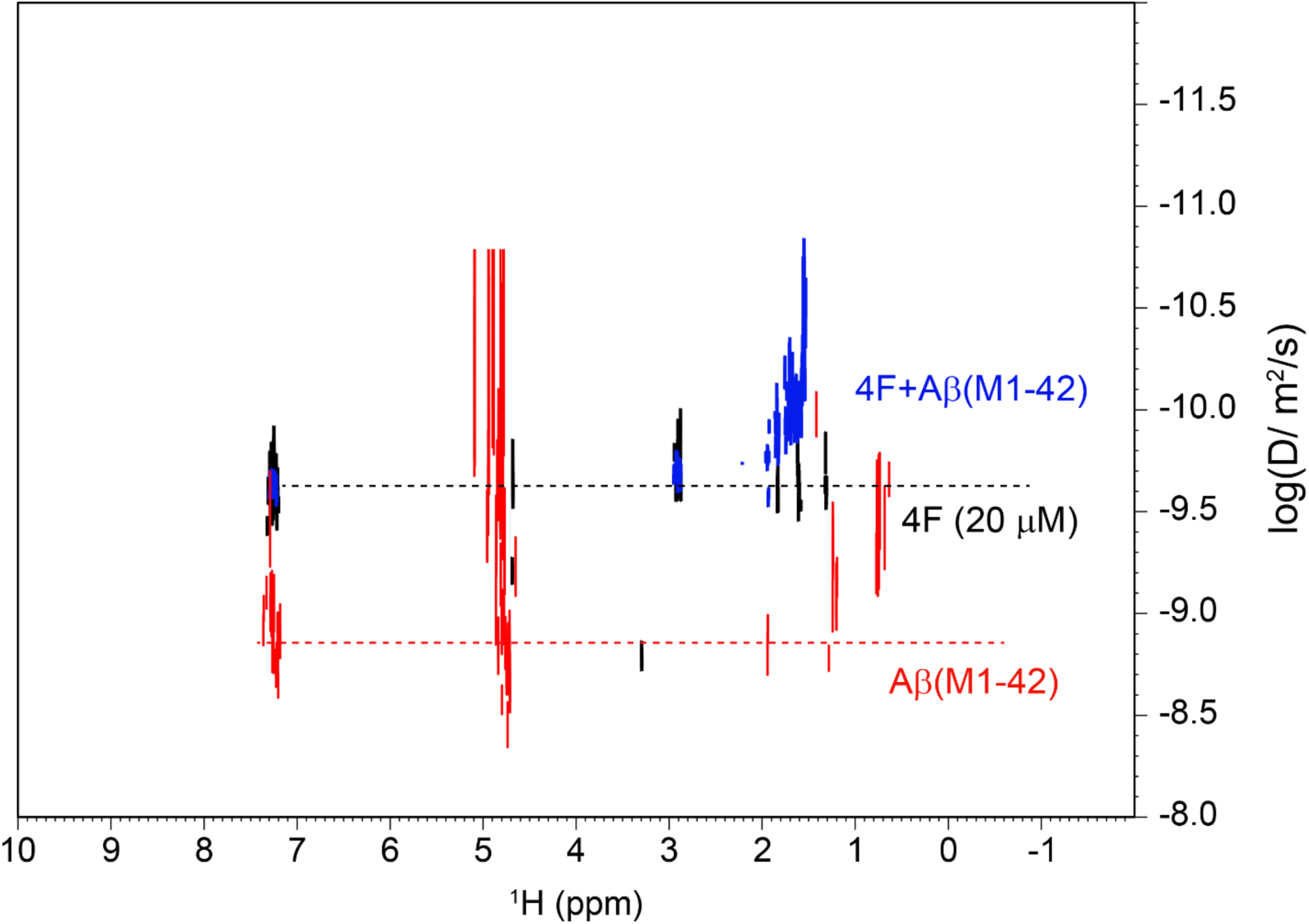
DOSY spectra of 20 µM 4F peptide in absence (black) and presence (blue) of equimolar Aβ (M1-42) in 100% D_2_O. The DOSY spectrum of 20 µM Aβ (M1-42) alone is shown in red. These data were obtained using a 500 MHz NMR spectrometer at 25 °C.

### Atomistic interaction between Aβ(M1-42) and 4F revealed using MD simulation

Next, we investigated the binding orientation of 4F peptide to Aβ(M1-42) using tryptophan fluorescence. The fluorescence emission of 5 µM 4F peptide (W2) was monitored for several days in presence and absence of equimolar Aβ(M1-42). In the absence of Aβ(M1-42), excitation of W2 residue in 4F at 295 nm showed fluorescent emission at λ_max_≈358 nm indicating its exposure to a polar environment (Fig. 5a). The 4F peptide mixed with equimolar concentration of Aβ(M1-42) showed an increase in W2 fluorescence with no significant spectral shift on day-1. However, differences were noticed on day-3 with a significant blue shift in W2 emission only in presence of Aβ(M1-42) indicating a non-polar environment of W2 in the Aβ-4F complex (Fig. 5a). Further incubation of these samples depicted disappearance of λ_max_ peak in Aβ-4F mixture on day-12 indicating the 4F peptides’ orientation in Aβ(M1-42) fibers exposing its W2 to the hydrophobic region of amyloid fibers. Such observations were previously observed for 4F peptide interacting with Aβ (1-40) indicating their preferential binding to Aβ peptides irrespective of their hydrophobicity.^28^

**Fig. 5.**
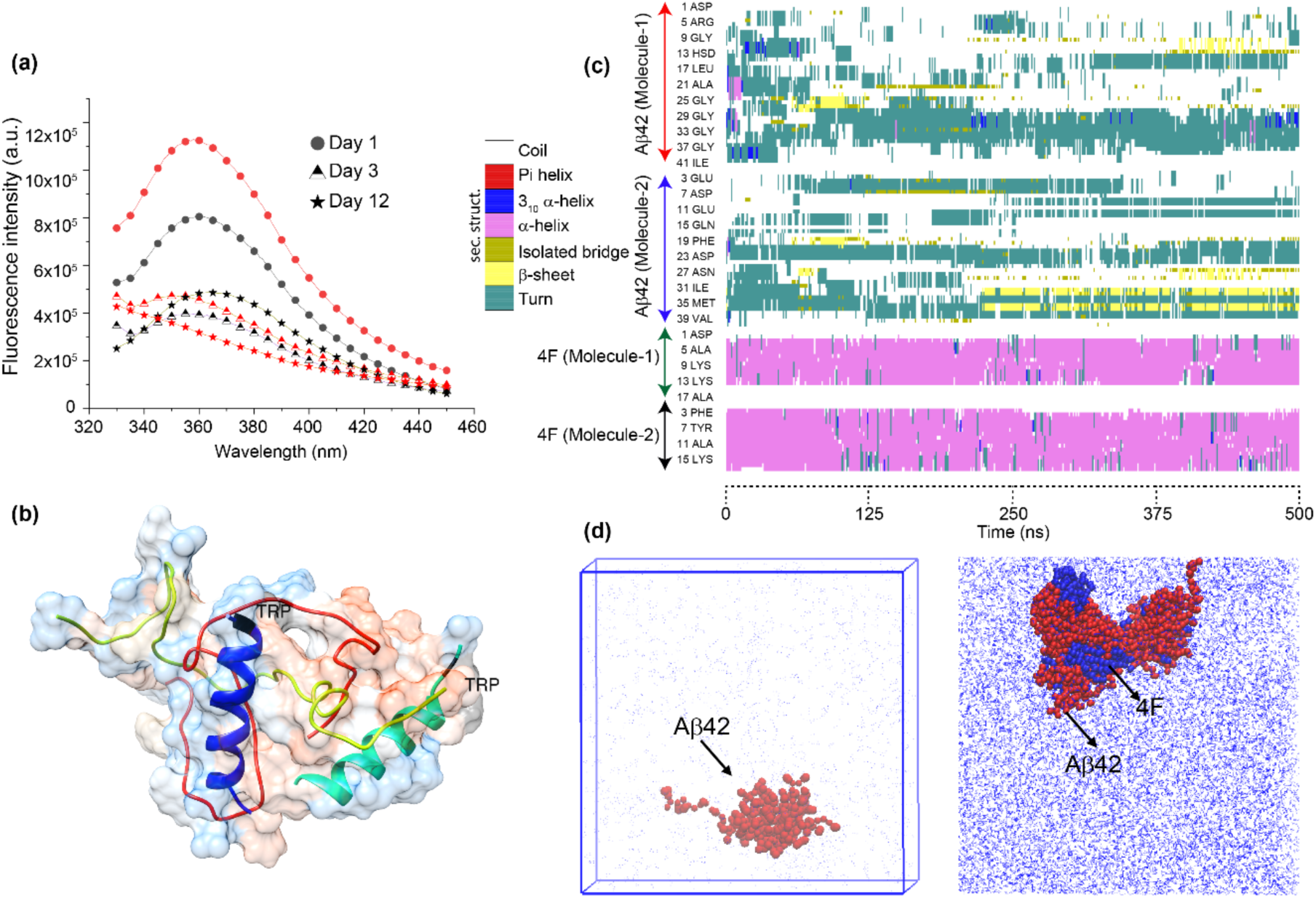
(a) Time-lapse tryptophan emission spectra of 5 µM 4F peptide in absence (black) or presence (red) of equimolar Aβ(M1-42) excited at 295 nm. The background fluorescence of the buffer are subtracted to correct spectra and are recorded in arbitrary units (a.u.). The samples are incubated at room temperature and fluorescence spectra are recorded for several days as indicated in different symbols at 25 °C. (b) MD snapshot illustrating interaction between 2 molecules of Aβ42 and 4F peptide (in cartoon) colored in yellow/red and blue/cyan, respectively. The molecular surface shows an increase in hydrophobicity from blue to orange. (c) The secondary structure evolution map of 4F-Aβ42 complex derived from 0.5 µs MD simulation as a function of amino acids. The secondary structure units are shown on the left (c). (d) Coarse-grained MD snapshots illustrating aggregation of 10 molecules Aβ42 (left) and hetero-aggregation complex of 4F-Aβ42 (right). The Aβ42 and 4F are shown in red and blue, respectively.

In an attempt to better understand the spatial orientation of 4F when bound to Aβ(1-42), we next performed hundreds of nanosecond all-atom molecular dynamics (MD) simulations. A significant secondary structural transition was identified in the partially folded Aβ(1-42) NMR structures (PDB ID: 1Z0Q) simulated in presence of 2 molecules of 4F (initially placed ∼ 0.5 nm away from Aβ) at the end of a 0.5 µs MD simulation. Both Aβ molecules depicted a random-coil conformation and were found to be tightly coupled with 4F with an ideal helix conformation (Fig. 5b). Secondary structure evolution map showed Aβ molecules mostly rendered random-coil or turns with a short transient β-sheet or 3_10_ α-helical structure when complexed with 4F. The atomistic structural model correlated well with CD and NMR observations described above that suggested a possible unstructured Aβ(M1-42) conformation when bound to 4F. In addition, the experimental observation of W2 spatial arrangement with an exposure to a non-polar environment was identified in the atomistic model of tetrameric structure that showed the W2 residues are buried inside the hydrophobic regions (Fig. 5b). The interacting residues in the 4F-Aβ(1-42) complex are listed in Table S1, which shows interaction sites that involve both N- and C-termini residues including central hydrophobic residues in Aβ(1-42). This result correlates with the average chemical shift perturbation obtained from ^15^N/^1^H SOFAST-HMQC NMR. Importantly MD calculations showed the formation of two salt-bridges between D7-K13 and K16-E16 in the 4F-Aβ(M1-42) complex. Several other interactions that include hydrogen bonds, alkyl and π-alkyl interactions are also identified in the 4F-Aβ(M1-42) complex and are listed in Table S1. Multi-microsecond MD simulation using coarse-grained models of 10 molecules of Aβ(1-42) distributed randomly showed an aggregated self-assembled complex at the end of 5 µs MD simulation (Fig. 5d, left). Interestingly, a random distribution of 10 molecules of Aβ(1-42) and 4F exhibited heterogenic assembly with 4F peptides buried inside Aβ(M1-42) oligomers (Fig. 5d, right). Notably, just as Aβ(1-42) self-assembling generates a single large particle, the spontaneous assembly of Aβ(1-42) and 4F produced a single heteromer complex (Fig. 5d) which correlates with the NMR observation of line-broadening and a uniform reduction in NMR signal intensity due to heteromerization (Fig. 4a and b). MM/PBSA approach estimated the free binding energy in the 4F-Aβ(M1-42) complex system. Although the MM/PBSA could not exactly imitate the experimental binding free energy, it could provide a comparative energy map by breaking down the free energy components that govern the complex formation. The 4F-Aβ(1-42) complex formation was found to be favored by Coulombic (ΔG_coul_), van der Waals (ΔG_vdw_) and non-polar solvation terms (ΔG_nps_), whereas the polar solvation energy (ΔG_ps_) opposed the complex formation (Table 1). These findings correlate well with the observed interacting residues between Aβ(1-42) and 4F contributing through salt-bridges and hydrogen bonds.

**Table 1.**
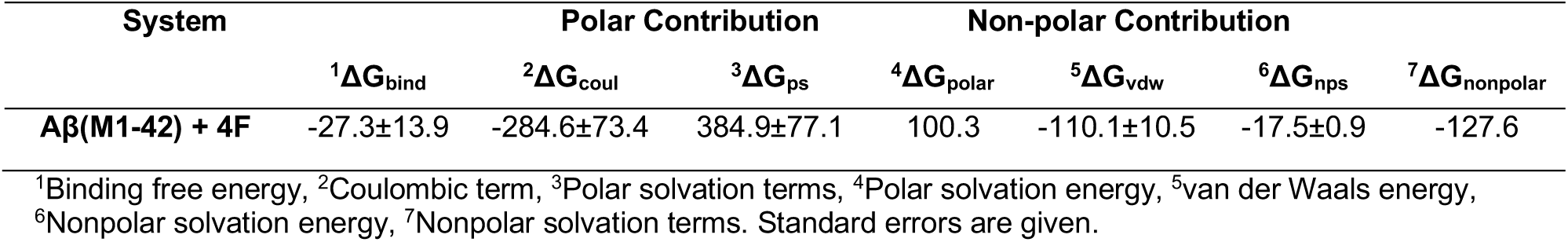
MM/PBSA based binding free energy (kcal mol^-1^) calculation.

### apoA-I mimetic peptide bound Aβ(M1-42) is neurotoxic

Controlling the neurotoxicity of Aβ(1-42) has remained a dominant research area in AD. Design and discovery of short peptide inhibitors for Aβ(1-42) are studied intensively. In this study, we also tested the impact of 4F peptide on the neurotoxicity of Aβ(M1-42) in differentiated SH-SY5Y neuroblastoma cells. Compared to the F-12 cell media, 4F peptide alone was found to be slightly but statistically insignificant neurotoxic (Fig. 6a) as previously observed for the full-length apoA-I.^15^ In contrast, high concentration 4F peptide increased cell proliferation in undifferentiated SH-SY5Y cells (Fig. S6). Although, a direct comparison of 4F’s cell toxicity on differentiated and undifferentiated cells is difficult, the differentiation of SH-SY5Y cells elevates 4F’s toxicity. A similar, differential phenotype between differentiated and undifferentiated rat neuronal cells treated with Aβ peptides has been reported.^46^ Remarkably, our results show that the Aβ(M1-42) complexed with 4F peptides exhibited higher neurotoxicity that Aβ alone. When 5 µM of Aβ(M1-42) monomers was used to treat differentiated SH-SY5Y cells, a cell viability of ≈75% was observed; and 5 µM of Aβ(M1-42) complexed with two or ten excess molar of 4F showed a reduction in viability of ≈65% and 45%, respectively (Fig. 6a). The MTT based cell-viability measurement on day-8 indicated that 4F generates toxic hetero-oligomers when complexed with Aβ(M1-42). Further fluorescence imaging of Aβ(M1-42) treated SH-SY5Y cells presented distinct morphological phenotypes in neurites in the presence and absence of 4F. The well-branched and connected SH-SY5Y cells observed on day-1 showed a gradual depletion in neurite outgrowth and connectivity treated with nocodazole, which is known to disrupt neurite growth and cause apoptosis (Fig. 6b and c). As compared to SH-SY5Y cell morphology observed in nocodazole and F-12 on Day-8, cells treated with 10 or 50 µM 4F peptide showed well-connected cells with more polarized soma cell bodies. In contrast to the MTT data, 4F mixed with Aβ(M1-42) did not significantly change neurite density, indicating either Aβ(M1-42)+4F induces changes in neurite morphology that are beyond the limits of detection in our assay or that cells are in the early stages of cytotoxic death. However, considering the MTT reduction assay that is commonly used to study Aβ toxicity in single cell cultures with high reproducibility,^47^ the observed results can be interpreted as 4F retards Aβ(M1-42) aggregation by generating a toxic hetero-oligomer species. This is in agreement with a previous finding that apolipoprotein fragments and Aβ(1-42) form toxic oligomers.

**Fig. 6.**
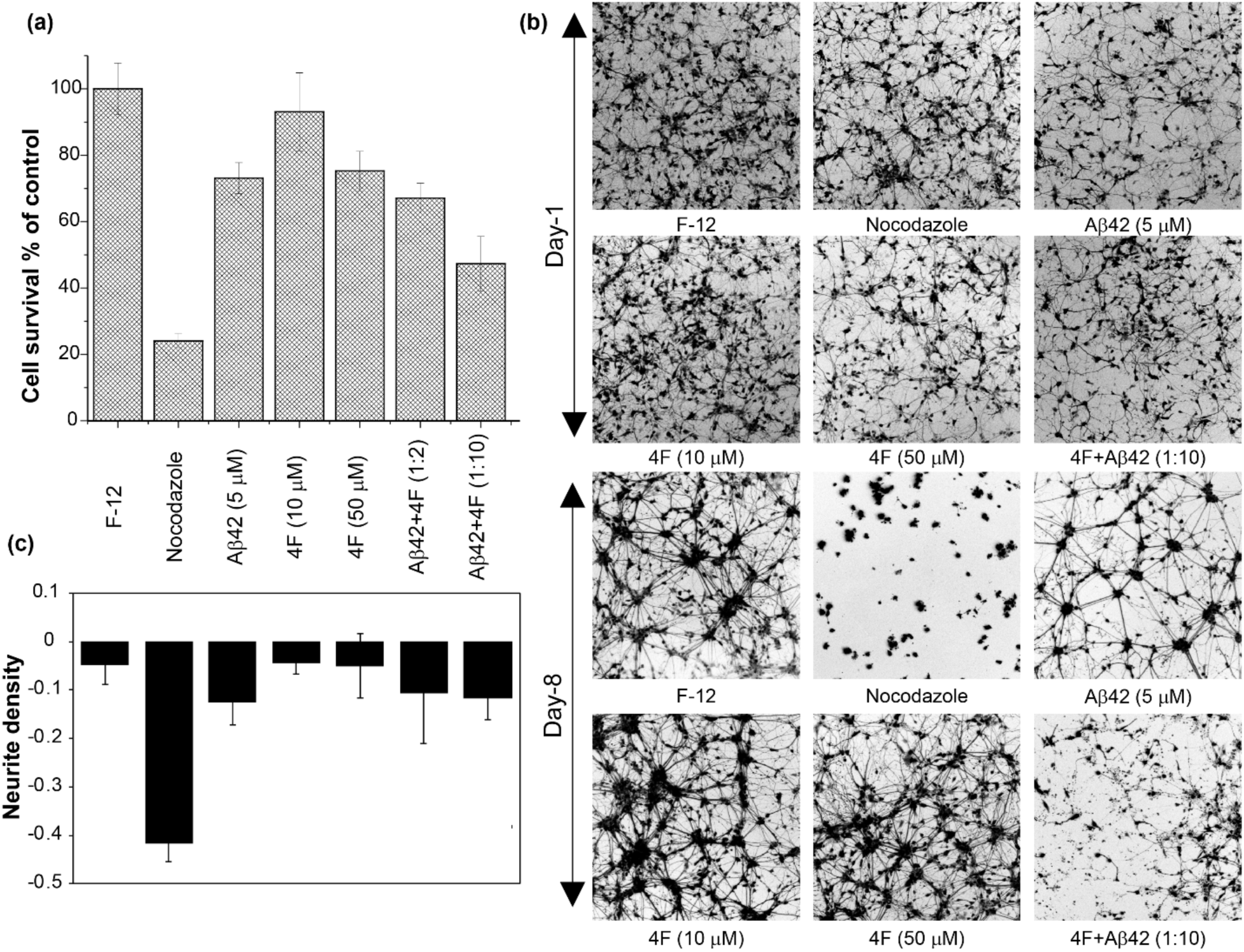
(a) Differentiated SH-SY5Y cell viability determined by MTT assay of 5 µM Aβ(M1-42)(denoted as Aβ42) incubated with 4F peptide at a variable concentration after 8 days. The cell-viability profiles are normalized using the % of live cells present in F-12 media without any additives. Shown are standard error from five replicates. (b) Fluorescence imaging of differentiated SH-SY5Y cells treated with Aβ (M1-42), 4F or mixture as described in Fig. 6A. Images were collected after treating cells on Day-1 and Day-8. Nocodazole is used as a positive control that shows significant damage in neurite growth. The cell-assay experiments were carried out by pre-incubating Aβ(M1-42) with 4F peptide at room temperature in 10 mM NaPi, pH 7.4. (c) Quantification of images from Fig.6b using *NeurphologyJ* to measure neurite density at each time point. Bars represent the average change in neurite density after treatment. Errors bars represent standard errors.

In summary, the apoA-I mimetic 4F peptide used in this study serves as an excellent model to explore the role of apolipoproteins or their fragments in modulating pathologically misfolded amyloidogenic Aβ(1-42) associated with AD. 4F shows a strong binding affinity for Aβ(M1-42) and is capable of retarding its aggregation. The hetero-oligomerization between 4F and Aβ(M1-42) generates water-soluble aggregates comprising of an unstructured Aβ(M1-42) and structured 4F that are found to be neurotoxic. While the 4F peptide was found to have protective role in proliferating cells, it’s association with amyloidogenic Aβ(M1-42) cause generation of cytotoxins. This finding strongly correlates between apolipoprotein fragmentations in brains thereby driving the disease progression. Taken together, the results based on our model study reported here urge further identification and characterization of natural apolipoprotein and its fragments in AD patients to aid in the development of chemical tools that can be used to reduce cellular toxicity.

## Experimental procedures

### Expression, purification and synthesis of peptides

The unlabeled Aβ(M1-42) (MDAEFRHDSGYEVHHQKLVFFAEDVGSNKGAIIGLMVGGVVIA) with an extra methionine at the N-terminus was recombinantly expressed in *E. coli* BL21 (DE3) pLysS Star in LB media as described elsewhere.^48^ The plasmid of Aβ(M1-42) was obtained from the Nowick laboratory at the University of California, Irvine, USA. Uniformly ^13^C and ^15^N isotope labelled Aβ(M1-42) was expressed by exchanging LB medium with M9 minimal medium containing ^15^NH_4_Cl and D-Glucose-^13^C_6_. We followed a published protocol to express ^15^N labelled Aβ(M1-42).^48^ The purified Aβ(M1-42) peptide powder (≈1 mg/mL) was treated with 5% NH_4_OH and aliquots of ≈0.1 mg/mL peptide were prepared after lyophilization. The lyophilized powder samples were re-suspended in the working buffer and vortexed for 15 s followed by sonication for 15 s in an ultrasonic bath sonicator. The small aggregates were next removed by centrifuging the sonicated peptide sample at 12,000 rpm for 5 minutes at 4 °C. The 4F peptide (Ac-DWFKAFYDKVAEKFKEAF-NH2) was synthesized by solid-phase from L-amino acids and purified by HPLC as described previously.^29,49^ 10 mM sodium phosphate (NaPi) buffer, pH 7.4 was used to dissolve Aβ(M1-42) and was used for all experiments (until and unless specified).

### Peptide aggregation assay using ThT fluorescence

The aggregation kinetics of 5 µM Aβ(M1-42) was monitored using 10 µM ThT fluorescence in the presence of varying concentrations of 4F peptide. A 96-well polystyrene plate was used with a sample volume of 100 µL/well in triplicate for all experiments. The aggregation kinetics were monitored using a microplate reader (Biotek Synergy 2) with slow shaking at 37 °C for 48 hours. The data were recorded at 3 minutes time interval with excitation and emission wavelengths at 440 and 485 nm, respectively. The averages of three replicates were plotted with respect to time.

### Transmission Electron Microscopy

TEM images of 5 µM Aβ(M1-42) incubated with 1 or 10 molar excess 4F peptide were taken using a HITACHI H-7650 transmission microscope (Hitachi, Tokyo, Japan) at 25 °C. The samples used for TEM analysis were collected from the 96-well plate at the end of 48 hours (peptides without the ThT dye). 20 µL/100 µL of sample volume were transferred to a collodion-coated copper grid and incubated for 5 minutes at room temperature followed by three rinses with 20 µL of double deionized water to remove buffer salts. The copper grid containing sample was next stained with 4 mL of 2% (w/v) uranyl acetate and incubated for 2 minutes followed by three rinses with 20 µL of double deionized water. The copper grid was next allowed to dry overnight under a vacuum desiccator at room temperature.

### Circular dichroism

Far-UV CD experiments were carried out for 20 µM Aβ(M1-42) dissolved in 10 mM NaPi in presence and absence of equimolar 4F at 25 °C using a JASCO (J820) spectropolarimeter. A 1 mm light-path length quartz cuvette containing 200 µL of sample was used for CD measurements. CD spectra of 20 µM of Aβ(M1-42) or 4F in absence of any additives were recorded and used as reference/control spectra. CD spectrum of 10 µM of Aβ(M1-42) titrated with 10% TFE was measured for comparative structural analysis. All samples were incubated at room temperature for time-lapse measurements.

### Tryptophan fluorescence

Tryptophan fluorescence was measured for 5 µM 4F peptide mixed with equimolar Aβ(M1-42) using a FluoroMax 4® from HoribaScientific® in continuous mode at 25 °C. The tryptophan was excited at 295 nm and the fluorescence emission was recorded from 330 to 450 nm (with a 5-nm bandwidth) with a delay time of 1 min per five scans using a 200-μL cuvette. The changes in fluorescence were monitored with respect to 4F peptide alone in the absence of Aβ(M1-42). All samples were incubated at room temperature for time-lapse measurements.

### NMR spectroscopy

All NMR samples were prepared in 10 mM NaPi buffer, pH 7.4 containing 10% deuterated water (v/v). Proton NMR spectra were acquired for unlabeled 20 µM Aβ(M1-42), 20 µM 4F and an equimolar (20 µM) mixture of Aβ(M1-42) and 4F using a 500 MHz NMR spectrometer at 25 °C. The NMR solution samples were next incubated under gentle agitation for 10 days before acquiring proton NMR spectra. The agitated 4F-Aβ(M1-42) mixed sample was next filtered using a 0.5 mL 30 kDa centrifugal filter (Amicon® Ultra-15) by adding 2M NaCl to remove electrostatically bound or free peptides from the solution. The washed buffer was collected, and tryptophan fluorescence was carried out to observe the presence of 4F. The filtered large aggregates/fibers were next treated with 2% deuterated SDS (v/v) and the sample was heated at 80 °C for 30 minutes to dissolve the fibers. Proton NMR spectra of the SDS treated sample was then acquired at 25 °C.

2D heteronuclear ^15^N/^1^H and ^13^C/^1^H (region of aromatic resonances) SOFAST-HMQC NMR experiments^50^ were carried out for uniformly labelled 20 µM Aβ(M1-42) in presence and absence of equimolar 4F in 10 mM NaPi, pH 7.4 containing 10% deuterated water (v/v). The 2D NMR experiments were recorded at 25 °C on a 800 MHz NMR spectrometer equipped with a 5 mm triple-resonance inverse detection TCI cryoprobe using 16 scans, 256 t1 increments and a 0.2 s recycle delay. For comparative structural study, the ^15^N/^1^H HMQC titration experiment was carried out for 20 µM Aβ(M1-42) dissolved in 10 mM NaPi, pH 7.4 containing 10% D_2_O and 10% deuterated 2,2,2 TFE (v/v). Diffusion Ordered SpectroscopY (DOSY) spectra were recorded using stimulated-echo with bipolar gradient pulses for diffusion with a gradient strength increment from 2 to 98% at 25 °C on a 500 MHz NMR spectrometer. NMR spectra were acquired with 16 gradient strength increments, 36,000 time domain data points in the t2 dimension, 3s recycle delay, and 100 ms diffusion delay. DOSY spectra were recorded for samples containing 20 µM Aβ(M1-42), 20 µM 4F or an equimolar mixture of both peptides (1:1) dissolved in 100% D_2_O.

### Cell viability assay

The cell-viability of human neuroblastoma (SH-SY5Y) cells was measured using MTT cell proliferation assay (Promega, G4000). SH-SY5Y cells were plated in a 96-well plate followed by differentiation in Neurobasal-A, GlutaMAX, B27, 1% penicillin/streptomycin, and 10 µM retinoic acid in a 5% CO_2_ humidified incubator at 37 °C. On day-5 of differentiation, SH-SY5Y cells were transduced with lentivirus encoding EGFP for neurite detection. Live-cells were imaged two days post-transduction prior to treatment (day-1) and after 8 days of (day-8) treatment on a Leica SP8 confocal microscope using a 10X objective to detect neurites. Differentiated SH-SY5Y cells (100 µL/well) were treated with 5 µM of Aβ(M1-42) incubated with or without a variable concentration of 4F (5, 10 and 50 µM). The effect of 4F peptide on undifferentiated SH-SY5Y cells was also measured at 5, 10 and 50 µM. MTT assay was performed on day-8 to measure the cell-viability following the manufacturer protocol for both differentiated and undifferentiated SH-SY5Y cells.

### MD simulations

All-atom and coarse-grained MD simulations were carried using GROMACS 5.0.7 ^51^ running parallel in SGI UV 3000 at the Institute for Protein Research, Osaka University, Japan. The 3D model of 4F peptide was built using an *ab-initio* modeling as described elsewhere ^28^. The NMR structure of Aβ42 (PDB ID: 1Z0Q) in aqueous solution was used as an initial model structure for MD simulation using charmm36 force-field ^52^. The MD systems were designed by placing two molecules of Aβ42 with 2 molecules of 4F peptide separated by a minimum distance of 0.5 nm in a box size of 9 nm x 9 nm x 9 nm. The simulation parameters were adopted from our previous studies.^28,36,53^ Briefly, all MD systems were neutralized and energy minimized using steepest-descent method followed by a short NVT and NPT equilibration MD. The equilibrated systems were next allowed for unrestrained MD simulation for a time-scale of 0.5 µs at 310.15 K. Coarse-grained MD simulations were carried out by randomly placing 10 Aβ42 molecules with or without 10 4F molecules using martini_v2.2P force field in a box size of 18 nm x 18 nm x 18 nm. MD trajectories were analyzed using Gromacs tools and graphical interpretations were done using VMD and Chimera. The GMXAPBS tool ^54,55^ was used to calculate the binding free energy from 500 MD snapshots retrieved at equal time interval from last 100 ns of 0.5 µs MD simulation. A detailed procedure of GMXAPBS calculation followed in this study is provided elsewhere.^55^

## Acknowledgements

This study was supported by funds from NIH (AG048934 to A.R.) and research in Y.W lab is supported by NIH grants (GM130331 and AG062225 to Y.W.). A part of this work (computational simulations) was performed under the International Collaborative Research Program of Institute for Protein Research, Osaka University, ICR-18-02.

## References

1 R. A. Moore, L. M. Taubner and S. A. Priola, Curr. Opin. Struct. Biol., 2009, 19, 14–22.

2 G. M. Ashraf, N. H. Greig, T. A. Khan, I. Hassan, S. Tabrez, S. Shakil, I. A. Sheikh, S. K. Zaidi, M. Akram, N. R. Jabir, C. K. Firoz, A. Naeem, I. M. Alhazza, G. A. Damanhouri and M. A. Kamal, CNS Neurol. Disord. Drug Targets, 2014, 13, 1280–93.

3 P. Sweeney, H. Park, M. Baumann, J. Dunlop, J. Frydman, R. Kopito, A. McCampbell, G. Leblanc, A. Venkateswaran, A. Nurmi and R. Hodgson, Transl. Neurodegener., 2017, 6, 6.

4 I. Benilova, E. Karran and B. De Strooper, Nat. Neurosci., 2012, 15, 349–357.

5 T. J. Esparza, H. Zhao, J. R. Cirrito, N. J. Cairns, R. J. Bateman, D. M. Holtzman and D. L. Brody, Ann. Neurol., 2013, 73, 104–119.

6 S. J. C. Lee, E. Nam, H. J. Lee, M. G. Savelieff and M. H. Lim, Chem. Soc. Rev., 2017, 46, 310–323.

7 L. Liao, D. Cheng, J. Wang, D. M. Duong, T. G. Losik, M. Gearing, H. D. Rees, J. J. Lah, A. I. Levey and J. Peng, J. Biol. Chem., 2004, 279, 37061–37068.

8 A. Mouchard, M. C. Boutonnet, C. Mazzocco, N. Biendon and N. Macrez, Sci. Rep., 2019, 9, 3989.

9 Y. Namba, M. Tomonaga, H. Kawasaki, E. Otomo and K. Ikeda, Brain Res., 1991, 541, 163–166.

10 P. L. Richey, S. L. Siedlak, M. A. Smith and G. Perry, Biochem. Biophys. Res. Commun., 1995, 208, 657–663.

11 T. Bilousova, M. Melnik, E. Miyoshi, B. L. Gonzalez, W. W. Poon, H. V. Vinters, C. A. Miller, M. M. Corrada, C. Kawas, A. Hatami, R. Albay, C. Glabe and K. H. Gylys, Am. J. Pathol., 2019, pii: S0002-9440(18)30596-0.

12 A. Kline, Alzheimer’s Res. Ther., 2012, 4.

13 P. B. Verghese, J. M. Castellano, K. Garai, Y. Wang, H. Jiang, A. Shah, G. Bu, C. Frieden and D. M. Holtzman, Proc. Natl. Acad. Sci., 2013, 110, E1807–E1816.

14 S. Ghosh, T. B. Sil, S. Dolai and K. Garai, 2019, 1–17.

15 A. C. Paula-Lima, M. A. Tricerri, J. Brito-Moreira, T. R. Bomfim, F. F. Oliveira, M. H. Magdesian, L. T. Grinberg, R. Panizzutti and S. T. Ferreira, Int. J. Biochem. Cell Biol., 2009, 41, 1361–1370.

16 J. Camacho, T. Moliné, A. Bonaterra-Pastra, S. Ramóny Cajal, E. Martínez-Sáez and M. Hernández-Guillamon, Front. Neurol., 2019, 10, 187.

17 T. Oda, P. Wals, H. H. Osterburg, S. A. Johnson, G. M. Pasinetti, T. E. Morgan, I. Rozovsky, W. B. Stine, S. W. Snyder, T. F. Holzman, G. A. Krafft and C. E. Finch, Exp. Neurol., 1995, 136, 22–31.

18 A. R. Nelson, A. P. Sagare and B. V. Zlokovic, Proc. Natl. Acad. Sci., 2017, 114, 8681–8682.

19 A. M. Wojtas, S. S. Kang, B. M. Olley, M. Gatherer, M. Shinohara, P. A. Lozano, C.-C. Liu, A. Kurti, K. E. Baker, D. W. Dickson, M. Yue, L. Petrucelli, G. Bu, R. O. Carare and J. D. Fryer, Proc. Natl. Acad. Sci., 2017, 114, E6962–E6971.

20 A. J. Doig and P. Derreumaux, Curr. Opin. Struct. Biol., 2015, 30, 50–56.

21 F. Belluti, A. Rampa, S. Gobbi and A. Bisi, Expert Opin. Ther. Pat., 2013, 23, 581–596.

22 S. A. Kotler, P. Walsh, J. R. Brender and A. Ramamoorthy, Chem. Soc. Rev., 2014, 43, 6692–6700.

23 M. M. M. Wilhelmus, R. M. W. De Waal and M. M. Verbeek, Mol. Neurobiol., 2007, 35, 203–216.

24 A. C. Kim, S. Lim and Y. K. Kim, Int. J. Mol. Sci., 2018, 19.

25 R. J. Baranello, K. L. Bharani, V. Padmaraju, N. Chopra, D. K. Lahiri, N. H. Greig, M. A. Pappolla and K. Sambamurti, Curr. Alzheimer Res., 2015, 12, 32–46.

26 W. Wang and X. Zhu, J. Neurochem., 2018, 147, 580–583.

27 D. Sawmiller, A. Habib, H. Hou, T. Mori, A. Fan, J. Tian, J. Zeng, B. Giunta, P. R. Sanberg, M. P. Mattson and J. Tan, Biol. Psychiatry, 2019, 86, 208–220.

28 B. R. Sahoo, T. Genjo, S. J. Cox, A. K. Stoddard, G. M. Anantharamaiah, C. Fierke and A. Ramamoorthy, J. Mol. Biol., 2018, 430, 4230–4244.

29 C. R. White, G. Datta, L. Wilson, M. N. Palgunachari and G. M. Anantharamaiah, Chem. Phys. Lipids, 2019, 219, 28–35.

30 C. B. Sherman, S. J. Peterson and W. H. Frishman, Cardiol. Rev., 2010, 18, 141–147.

31 L. T. Bloedon, R. Dunbar, D. Duffy, P. Pinell-Salles, R. Norris, B. J. DeGroot, R. Movva, M. Navab, A. M. Fogelman and D. J. Rader, J. Lipid Res., 2008, 49, 1344–1352.

32 F. Su, K. R. Kozak, S. Imaizumi, F. Gao, M. W. Amneus, V. Grijalva, C. Ng, A. Wagner, G. Hough, G. Farias-Eisner, G. M. Anantharamaiah, B. J. Van Lenten, M. Navab, A. M. Fogelman, S. T. Reddy and R. Farias-Eisner, Proc. Natl. Acad. Sci., 2010, 107, 19997–20002.

33 C. Nerelius, A. Sandegren, H. Sargsyan, R. Raunak, H. Leijonmarck, U. Chatterjee, A. Fisahn, S. Imarisio, D. A. Lomas, D. C. Crowther, R. Stromberg and J. Johansson, Proc. Natl. Acad. Sci., 2009, 106, 9191–9196.

34 H. Levine III, Amyloid, 2007, 14, 185–197.

35 F. E. Cohen and J. W. Kelly, Nature, 2003, 426, 905–909.

36 B. R. Sahoo, T. Genjo, T. W. Nakayama, A. K. Stoddard, T. Ando, K. Yasuhara, C. A. Fierke and A. Ramamoorthy, Chem. Sci., 2019, 10, 3976–3986.

37 R. Limbocker, S. Chia, F. S. Ruggeri, M. Perni, R. Cascella, G. T. Heller, G. Meisl, B. Mannini, J. Habchi, T. C. T. Michaels, P. K. Challa, M. Ahn, S. T. Casford, N. Fernando, C. K. Xu, N. D. Kloss, S. I. A. Cohen, J. R. Kumita, C. Cecchi, M. Zasloff, S. Linse, T. P. J. Knowles, F. Chiti, M. Vendruscolo and C. M. Dobson, Nat. Commun., 2019, 10, 225.

38 B. R. Sahoo, T. Genjo, M. Bekier, S. J. Cox, A. K. Stoddard, M. Ivanova, K. Yasuhara, C. A. Fierke, Y. Wang and A. Ramamoorthy, Chem. Commun. (Camb)., 2018, 54, 12883–12886.

39 T. L.M., B. T., J. L., R. S.K., Y. K.L., Y. C., P. W.W., C. L.B., M. C.A., V. H.V., V. E. L.J., F. D.W., E. S., B. G., G. K.H. and L. M.J., J. Biol. Chem., 2013, 288, 5914–5926.

40 K. Garai, P. B. Verghese, B. Baban, D. M. Holtzman and C. Frieden, Biochemistry, 2014, 53, 6323–6331.

41 G. S. Getz and C. A. Reardon, J. Inflamm. Res., 2011, 4, 83–92.

42 M. Serra-Batiste, M. Ninot-Pedrosa, M. Bayoumi, M. Gairí, G. Maglia and N. Carulla, Proc. Natl. Acad. Sci., 2016, 113, 10866–10871.

43 C. Dammers, L. Gremer, K. Reiß, A. N. Klein, P. Neudecker, R. Hartmann, N. Sun, H. U. Demuth, M. Schwarten and D. Willbold, PLoS One, 2015, 10, e0143647.

44 J. Roche, Y. Shen, J. H. Lee, J. Ying and A. Bax, Biochemistry, 2016, 55, 762–775.

45 T. Asakura, K. Taoka, M. Demura and M. P. Williamson, J. Biomol. NMR, 1995, 6, 227–236.

46 T. Liu, G. Perry, H. W. Chan, G. Verdile, R. N. Martins, M. A. Smith and C. S. Atwood, J. Neurochem., 2004, 88, 554–563.

47 R. Rönicke, A. Klemm, J. Meinhardt, U. H. Schröder, M. Fändrich and K. G. Reymann, PLoS One, 2008, 9, e3236.

48 S. Yoo, S. Zhang, A. G. Kreutzer and J. S. Nowick, Biochemistry, 2018, 57, 3861–3866.

49 G. Datta, M. Chaddha, S. Hama, M. Navab, A. M. Fogelman, D. W. Garber, V. K. Mishra, R. M. Epand, R. F. Epand, S. Lund-Katz, M. C. Phillips, J. P. Segrest and G. M. Anantharamaiah, J. Lipid Res., 2001, 42, 1096–104.

50 P. Schmieder,

51 D. Van Der Spoel, E. Lindahl, B. Hess, G. Groenhof, A. E. Mark and H. J. C. Berendsen, J. Comput. Chem., 2005, 26, 1701–1718.

52 J. Huang and A. D. Mackerell, J. Comput. Chem., 2013, 34, 2135–2145.

53 B. R. Sahoo, J. Maharana, G. K. Bhoi, S. K. Lenka, M. C. Patra, M. R. Dikhit, P. K. Dubey, S. K. Pradhan and B. K. Behera, Mol. BioSyst., 2014, 10, 1104–1116.

54 D. Spiliotopoulos, A. Spitaleri and G. Musco, PLoS One, 2012, 7, e46902.

55 B. R. Sahoo, J. Maharana, M. C. Patra, G. K. Bhoi, S. K. Lenka, P. K. Dubey, S. Goyal, B. Dehury and S. K. Pradhan, Colloids Surfaces B Biointerfaces, 2014, 121, 307–318.

