## Supporting information for "Apolipoprotein A-I Mimetic 4F Peptide Generates Amyloid Cytotoxins by Forming Hetero-oligomers with β-amyloid"

**Table S1**. List of Aβ(1-42) residues interacting with 4F peptide obtained from 0.5 μs MD simulation.

| **Aβ42-4F complex** | | |
|---|---|---|
| C:LYS13:HZ3 | **B:**ASP7:OD2 | Salt Bridge; Attractive Charge |
| **B:**LYS16:HZ1 | C:GLU16:OE1 | Salt Bridge; Attractive Charge |
| C:LYS9:NZ | **B:**ASP23:OD1 | Attractive Charge |
| C:PHE18:HN | **B:**HSD13:O | Conventional Hydrogen Bond |
| D:TYR7:HH | **B:**GLY33:O | Conventional Hydrogen Bond |
| **A:**HSD14:HD1 | **B:**ASP23:OD2 | Conventional Hydrogen Bond |
| **A:**GLN15:HN | **B:**GLU22:O | Conventional Hydrogen Bond |
| **A:**LYS16:HN | **B:**GLU22:O | Conventional Hydrogen Bond |
| **A:**PHE20:HN | **B:**ALA2:O | Conventional Hydrogen Bond |
| **A:**SER26:HN | **B:**VAL18:O | Conventional Hydrogen Bond |
| **A:**ASN27:HD21 | **B:**GLY38:O | Conventional Hydrogen Bond |
| **A:**LYS28:HZ2 | **B:**LYS16:O | Conventional Hydrogen Bond |
| **A:**LYS28:HZ3 | **B:**ALA42:O | Conventional Hydrogen Bond |
| **A:**GLY29:HN | **B:**GLY37:O | Conventional Hydrogen Bond |
| **A:**ALA30:HN | **B:**GLY37:O | Conventional Hydrogen Bond |
| **B:**PHE4:HN | **A:**VAL18:O | Conventional Hydrogen Bond |
| **B:**GLU11:HN | C:LYS15:O | Conventional Hydrogen Bond |
| **B:**VAL12:HN | C:GLU16:O | Conventional Hydrogen Bond |
| **B:**GLN15:HE21 | C:PHE18:O | Conventional Hydrogen Bond |
| C:ALA17:HA | **B:**VAL12:O | Carbon Hydrogen Bond |
| **A:**HSD13:HA | **B:**ALA21:O | Carbon Hydrogen Bond |
| **A:**HSD14:HA | **B:**GLU22:O | Carbon Hydrogen Bond |
| **A:**PHE19:HA | **B:**ALA2:O | Carbon Hydrogen Bond |
| **A:**GLY25:HA1 | **B:**VAL18:O | Carbon Hydrogen Bond |
| **A:**GLY29:HA1 | **B:**VAL39:O | Carbon Hydrogen Bond |
| **B:**GLU3:HA | **A:**VAL18:O | Carbon Hydrogen Bond |
| **B:**GLU22:HA | **A:**LYS16:O | Carbon Hydrogen Bond |
| **B:**ASP23:HA | **A:**HSD13:O | Carbon Hydrogen Bond |
| C:PHE14 | D:PHE18 | Pi-Pi T-shaped |
| **B:**GLY33:C,O;LEU34:N | D:PHE3 | Amide-Pi Stacked |
| C:ALA5 | **B:**VAL24 | Alkyl |
| C:LYS13 | **B:**LYS16 | Alkyl |
| D:VAL10 | **A:**ILE32 | Alkyl |
| D:VAL10 | **A:**VAL40 | Alkyl |
| D:VAL10 | **B:**VAL39 | Alkyl |
| D:ALA11 | **B:**ILE41 | Alkyl |
| D:LYS13 | **A:**ILE32 | Alkyl |
| D:ALA17 | **A:**VAL36 | Alkyl |
| **B:**ALA21 | **A:**VAL12 | Alkyl |
| **B:**ALA21 | **A:**LYS16 | Alkyl |
| C:PHE6 | **B:**VAL18 | Pi-Alkyl |
| C:TYR7 | **A:**ILE31 | Pi-Alkyl |
| C:TYR7 | **A:**MET35 | Pi-Alkyl |
| D:PHE3 | **A:**ALA42 | Pi-Alkyl |
| D:PHE6 | **A:**VAL40 | Pi-Alkyl |
| D:PHE6 | **A:**ALA42 | Pi-Alkyl |
| D:PHE6 | **B:**VAL39 | Pi-Alkyl |
| D:PHE14 | **A:**LYS28 | Pi-Alkyl |
| D:PHE14 | **B:**ILE41 | Pi-Alkyl |
| **A:**PHE19 | **B:**ALA2 | Pi-Alkyl |
| **A:**PHE20 | **B:**ALA2 | Pi-Alkyl |
| **B:**PHE4 | **A:**ALA21 | Pi-Alkyl |
| **B:**PHE19 | **A:**VAL18 | Pi-Alkyl |

**[Chain A: and B: for the Aβ(1-42) peptide; Chain C: and D: for the 4F peptide]**

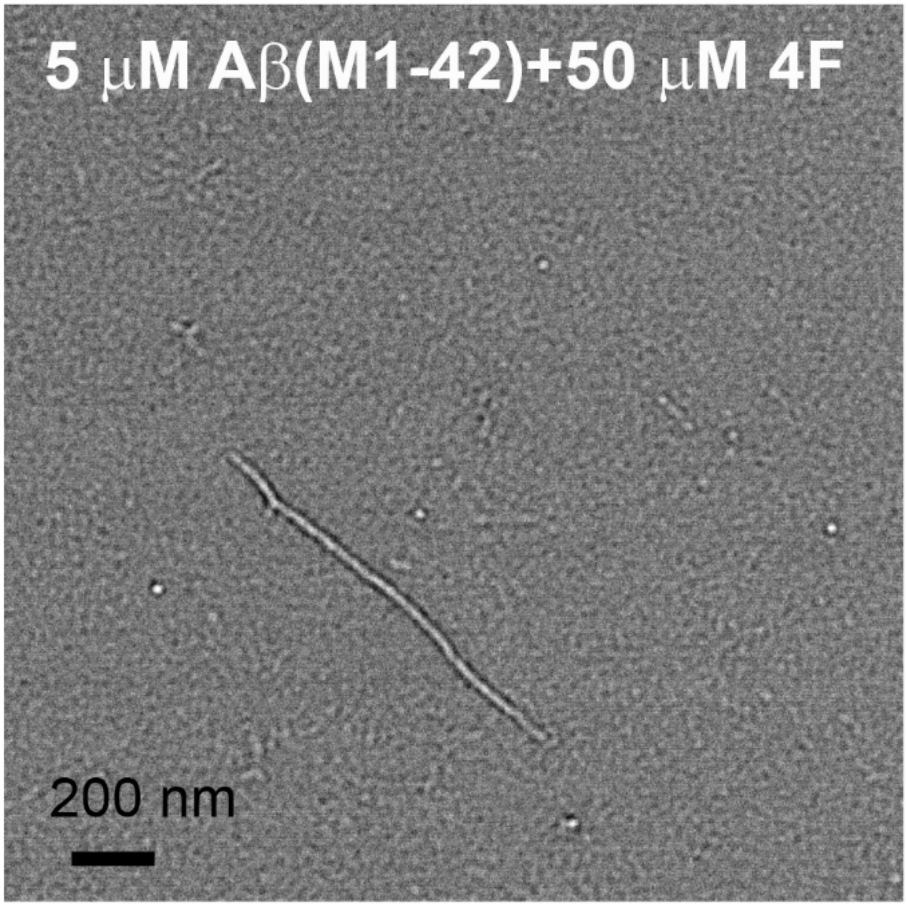


**Figure S1.** Transmission electron microscopy (TEM) image showing a straight fibrillary morphology of 5 μM Aβ(M1-42) incubated with 50 μM 4F peptide.

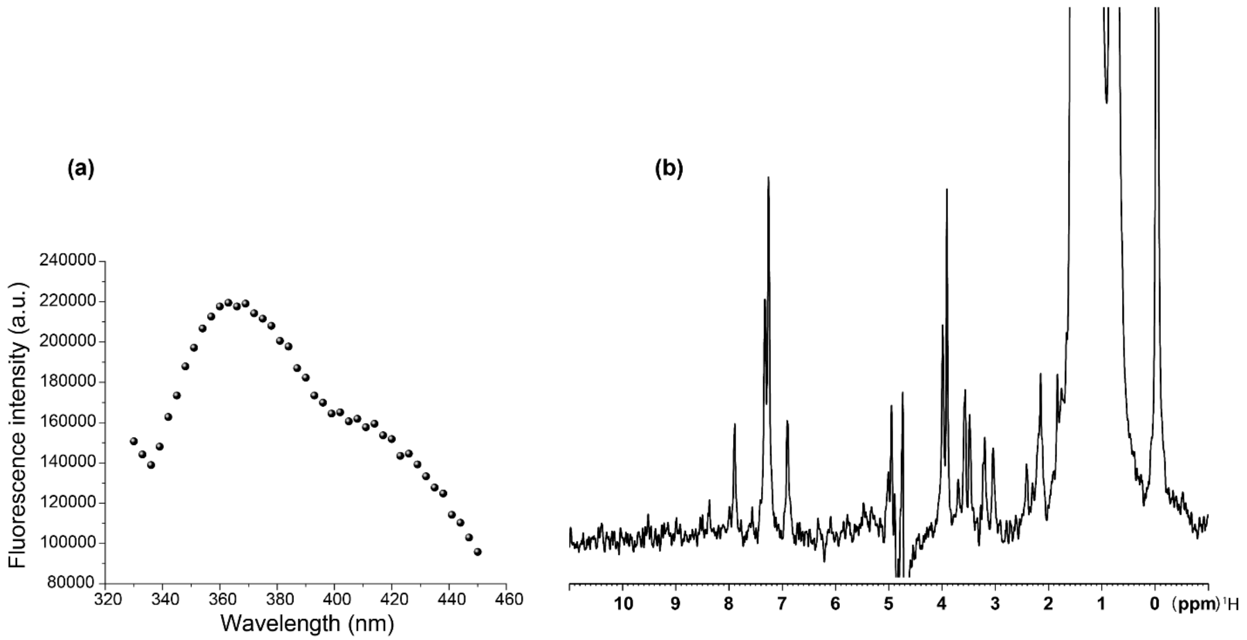


**Figure S2.** (a) Tryptophan fluorescence of sample waste collected from 4F-Aβ(M1-42) after day-10. The sample waste was collected by washing 4F-Aβ42(M1-42) mixed sample in 2M NaCl two times using a 30 kDa centrifugal filter (see methods). (b) $^{1}H$ NMR spectrum of a 2% deuterated-SDS treated filtered sample (see the Methods Section in the main text) recorded on a 500 MHz NMR spectrometer at 25 ℃.

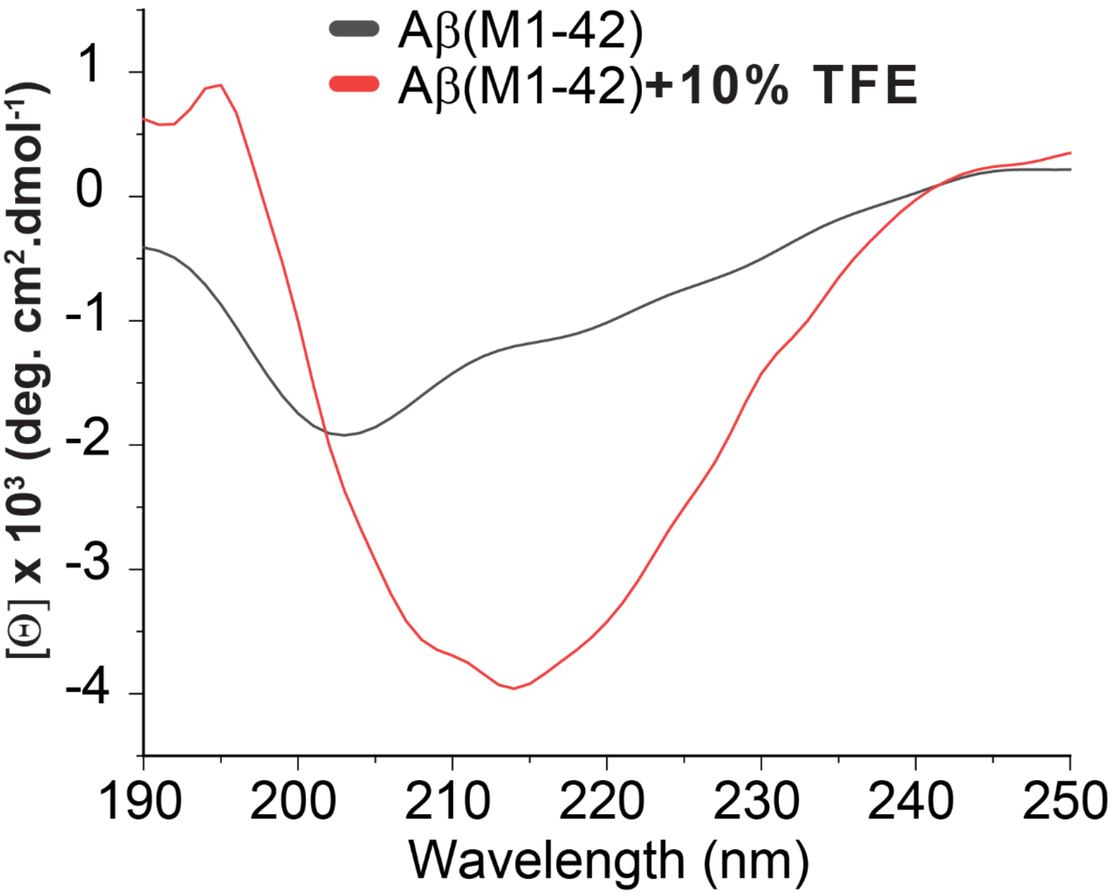


**Figure S3.** CD spectra showing a conformational change in Aβ(M1-42) peptide: 20 μM Aβ(M1-42) in absence (black) and presence (red) of 10% TFE.

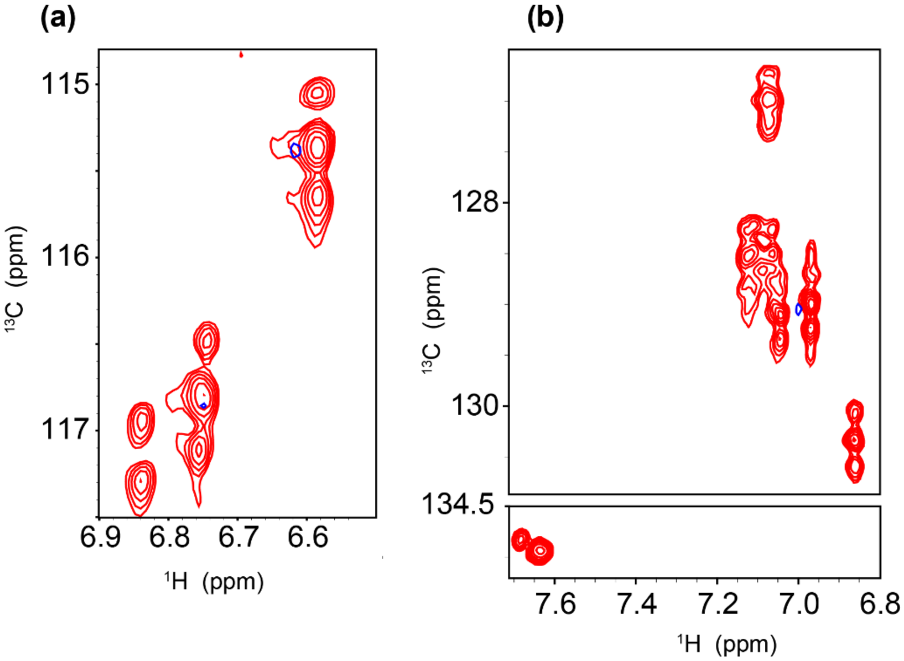


**Figure S4.** Aromatic regions of 2D $^{1}H/^{13}C$ SOFAST-HMQC NMR spectra of 20 μM Aβ(M1-42) in absence (red) and presence (blue) of an equimolar 4F peptide. The observed $^{1}H/^{13}C$ resonances are from the aromatic side chains of phenylalanine and tyrosine residues (Y10/F19/F20) of the Aβ(M1-42) peptide.

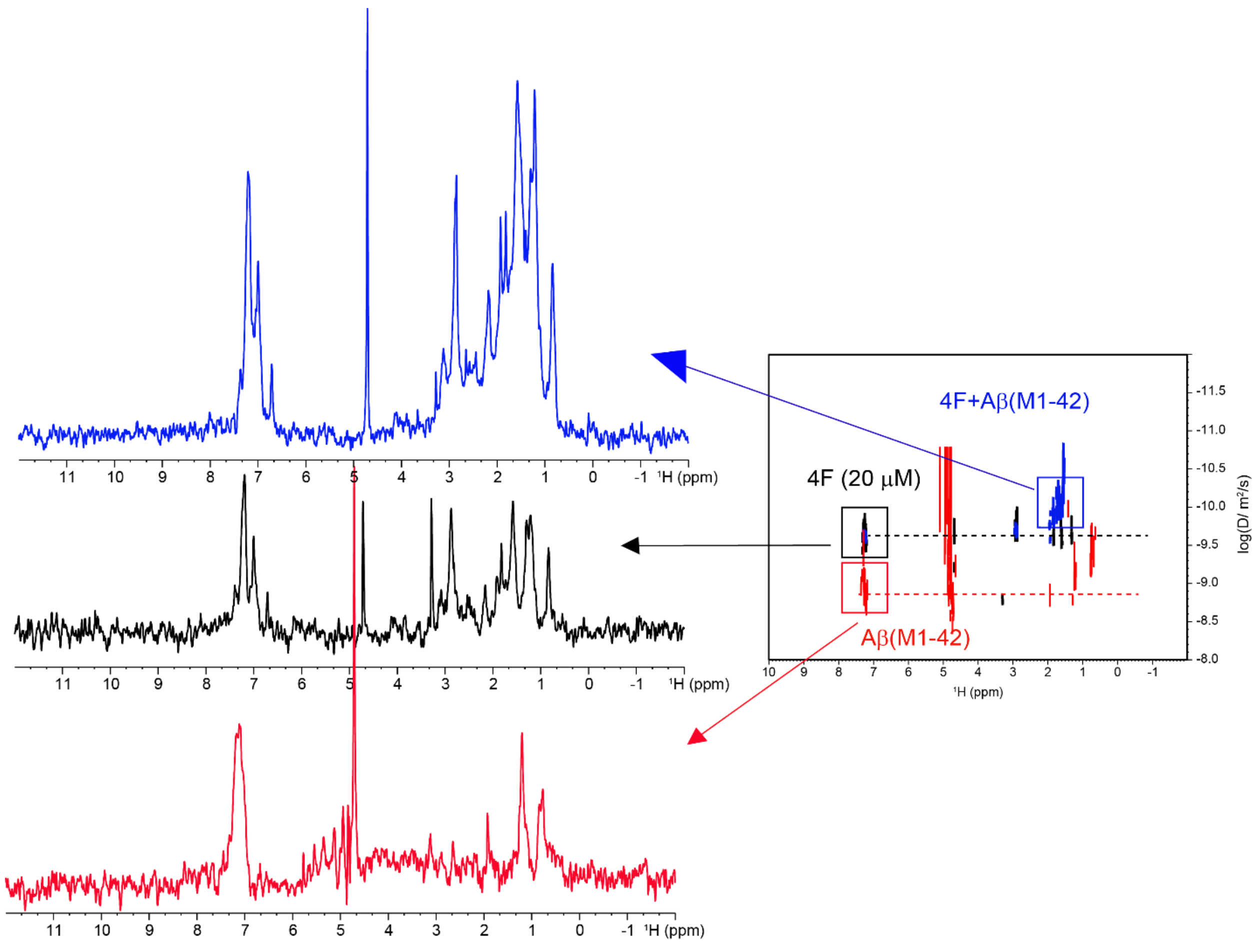


**Figure S5.** 1D spectra of 20 μM 4F peptide in absence (black) and presence (blue) of equimolar Aβ(M1-42) and 20 μM of Aβ(M1-42) in absence of 4F (red) in 100% $D_2O$ extracted from the 2D DOSY spectrum shown in right (also shown in Fig.4 in the main text). The DOSY spectra were obtained using a 500 MHz NMR spectrometer at 25 ℃.

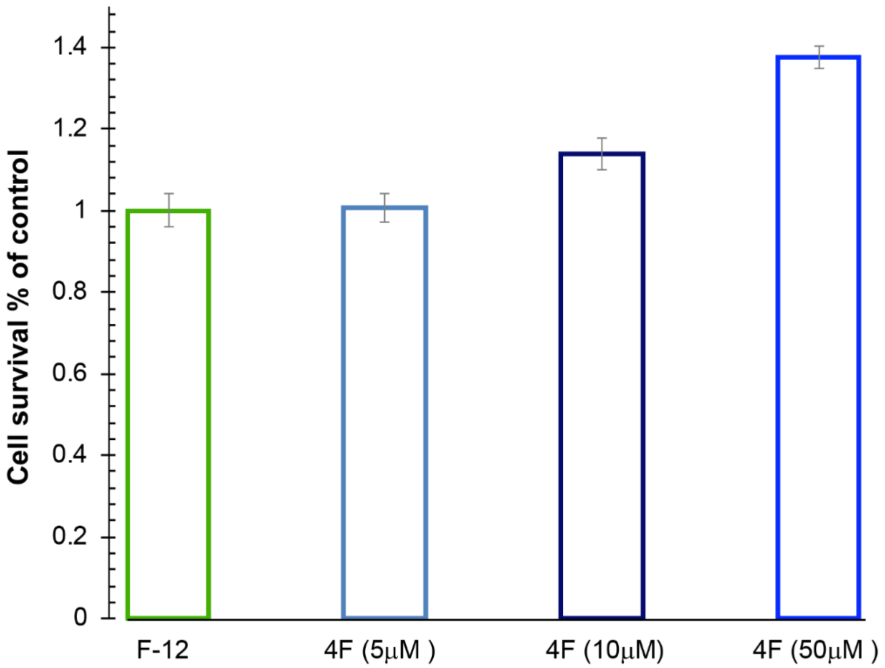


**Figure S6.** Effect of 4F peptide on the cell viability of undifferentiated SH-SY5Y cells as measured using MTT reagent. The cell-viability profiles are normalized using the % of live cells present in F-12 media without any additives.
